# Reckless driving: Improved phosphorous availability is a most likely important driver and species killer in angiosperm evolution

**DOI:** 10.1101/687939

**Authors:** Stefan Olsson

## Abstract

Plants show a large range of genome sizes with a more than thousand fold difference between the largest and the smallest genomes (Bennett, M. D., 1972; Bennett, 2005; Knight, 2005; Gregory et al., 2007; Bai et al., 2012; Kelly et al., 2015). Since, 1. DNA phosphorous is a phosphorous pool not possible to recycle within the cell. 2. Carnivorous plants with very small genomes living in low phosphorous environments uses carnivory to supply phosphorous (Leushkin et al., 2013). 3. The genus *Fritillaria* with very large genomes is dependent on mycorrhiza (Wang & Qiu, 2006; Kelly et al., 2015) probably mainly for supply of P, I decided to investigate if lifestyles connected with releases from P-limitation correlate with 1C-genome sizes. For the investigation I used available lifestyle data from several websites and articles (Wang & Qiu, 2006; Lysak et al., 2008) together with the Angiosperm C-values available at Kew http://data.kew.org/cvalues/. Based on this analysis I suggest the following order of P restriction for genome size expansions, from the least restricted to the most restricted, for the different lifestyles: Myco-heterotrophs<Parasitic plants<Mycorrhizal plants<Non-Mycorrhizal plants<Carnivorous plants. The data further suggest that a general improved P status most likely precedes whole genome duplication events and that uneven ploidy increases are more punishing than even ploidy most probably since uneven ploidy increases cannot be followed by adaptive recombination/selection. Thus P-limitation is a likely main restraint for angiosperm evolution and genome-expansion. However, relaxing this restraint leads plants into an evolutionary dead end with large nuclei (Bennetzen & Kellogg, 1997) and high P demand.

## MAIN

Phosphorous is a common limiting nutrient for plant growth (Lambers & Plaxton, 2015). The, compared to other element, relatively large proportion of non-recyclable P in DNA that cannot be lowered in response to nutrient limitations in phytoplankton cells (Berdalet, Latasa & Estrada, 1994). This should restrain genome expansions in eukaryotic cells towards unnecessary large genomes. since this would limit growth and survival in natural P nutrient limiting environments since under P-limiting conditions RNA necessary for protein synthesis and growth works as a P-reservoir (Berdalet, Latasa & Estrada, 1994). A small genome should thus be advantageous under P limiting conditions. However, in plants, gene duplications and especially whole genome duplication events followed by evolving new uses for the duplicated genes are often seen as fast forward mechanism for plant evolution. Such duplications are generally followed by fast removal of duplicated genes not finding relevant functions in double copies or as new genes and the plant revert to the diploid state although traces of the gene duplication event is left in the genome (Leitch, I. J. & Bennett, M. D., 2004; Šmarda et al., 2013; Han et al., 2018). There is also no correlation between size of genome and phylogenetic history in severely nutrient limited karst plants (Kang, Wang & Huang, 2015) even if phosphorous limitations after whole genome expansion events can restrain plants with large genomes and limit their competitiveness (Šmarda et al., 2013). However, accumulation of non-coding repeats seems to be a more common cause of genome expansion (Kelly et al., 2015), and contraction if there is a severe P-limitation (Leushkin et al., 2013; Šmarda et al., 2013). The very small genomes of carnivorous plants have normal amounts of functional genes as well as regulatory elements between genes and seem to have mechanisms actively restricting genome expansion through accumulation of repeats (Leushkin et al., 2013). Similarly, whole genome duplication events seem to have been followed by genome reductions (most probably by removing non-coding repeats) since ploidy level is inversely correlated with size of the basic genome (Leitch, I. J. & Bennett, M. D., 2004).

Hypotheses: 1. Niche phosphorous availability constrains plant genome sizes. 2. Freed from this constrain genome expands by two processes, gene duplications and probably more frequently by accumulation of non-coding repeats. 3. Phosphorous constraints mainly favor genome contraction by removal of repeats and but also by removal of duplicated genes that has not found new functions.

If these hypotheses are supported plants with very large genomes should occupy niches with relaxed phosphorous limitations while plants having very small genomes should occupy niches where there are strict phosphorous limitations. There should be little correlation between genome size and evolutionary history since accumulation of non-coding repeats does not lead to speciation but a strong correlation between plant phosphorous content and genome size.

It is not practically possible to experimentally test drivers (causes) of evolutionary changes since that would take thousands or millions of years. But it is possible from the literature to come up with hypotheses that can be tested in already available data if that data can be re-arranged and compiled in new ways guided by the hypotheses without post-hoc hypothesis testing.

First, I collected data on different lifestyles that could be connected to different phosphorous availability. Carnivorous plants live in very phosphorous poor environments and are known to use the caught small animals to supply phosphorous (Leushkin et al., 2013). One of the main roles of the mycorrhizal symbiosis is to supply phosphorous to the plant (Smith, Anderson & Smith, 2015) and finally, plants that are parasitic on other plants as well as plants like the more or less mycoheterothrophic orchids could be expected to be least restricted in P availability. List of plant species and their genome sizes (1C values) belonging to the 5 different groups were produced from the relevant references and compared (see Methods and Supplementary files **Data S1-4**).

The data presented in **Fig. 1** lends support to the hypothesis that genome size correlates well with lifestyles relaxed in P limitation. In addition, the sorted percentile plot (**Fig. 1B**) shows the overlap in genome sizes between groups and also highlights the comparably low genome size for some of the parasitic plants (to the left in **Fig. 1B**). These turns out to be mainly hemiparasites (**Fig. 1B**) and grow independently as seedlings giving further support to the general hypothesis.

**Figure 1.**
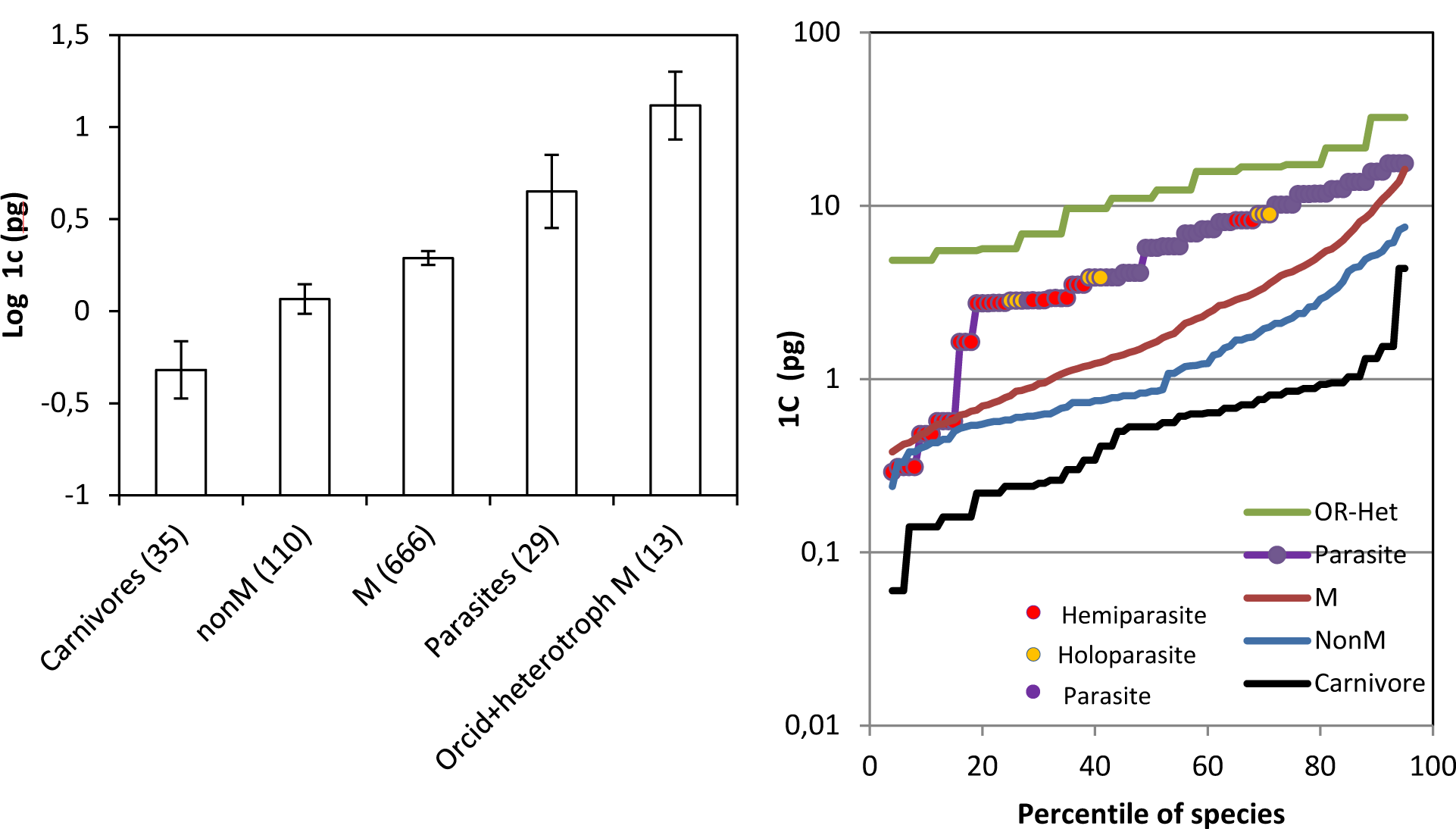
Log genome size in plants with different lifestyles Carnivores (Carnivore), non-Mycorrhizal (nonM) Mycorrhizal (M), Parasites in other plants (Parasite) and Orchids-heterotrophic mycorrhizal (OR-Het). Numbers in brackets represents number of plant species co-existing on lists for respective lifestyles and in the genome size database. A. Mean values and confidence intervals. Error bars represents 95% confidence intervals calculated using non-parametric methods to avoid assuming normal distribution. Mean values with non-overlapping confidence intervals are significantly different (p=≪0.05). B. Sorted percentile plots from small genome size to high genome sizes.

The phosphorous content of the seeds in combination with phosphorous demand of the plant is a determinant for the success of plant establishment (White & Veneklaas, 2012). A perennial lifestyle can relax the demand for phosphorous since there is less need for producing phosphorus containing seeds after only one season. Thus, being perennial could potentially offset higher phosphorous demand due to a large genome. Previously it has been noted that perennials have larger genome sizes in general (Bennett, M. D., 1972) and information on perennial vs annual is also available in the Kew database on 1c sizes. I could then compare the genome sizes for annuals vs perennials but also decided to do so for non-mycorrhizal annuals/perennials, mycorrhizal annuals/perennials, and the strictly non-mycorrhizal family Brassicaceae annuals/perennials (**Fig. 2A**). The previous observation (Bennett, M. D., 1972) that perennials in general have larger genomes compared to annuals was duplicated. There was however no significant difference between non-mycorrhizal plants annual vs perennial or between mycorrhizal plants annual vs perennial. This could be the result of a low sample number since only the plant species that could be included in the analysis was the ones appearing on both the mycorrhizal plant list (Wang & Qiu, 2006) and the list of annual and perennials with known 1C values (from the Kew database). The family Brassicaceae contains both annual and perennial plants and is not known to form any type of mycorrhizal symbioses so the difference in genome size connected with annual and perennial lifestyles was investigated in this family and included in **Fig. 2A** and in a **Fig. 2B**. Interestingly, the perennial Brassicaceae species with the smallest genomes are mainly endemic to the geologically young Canary Islands (Goodson, Santos-Guerra & Jansen, 2006; Goodson, 2007) and belonging to genera that are otherwise mostly annual. This could possibly point to a recent shift to a perennial lifestyle (Supplementary file **Data S1**).

**Figure 2.**
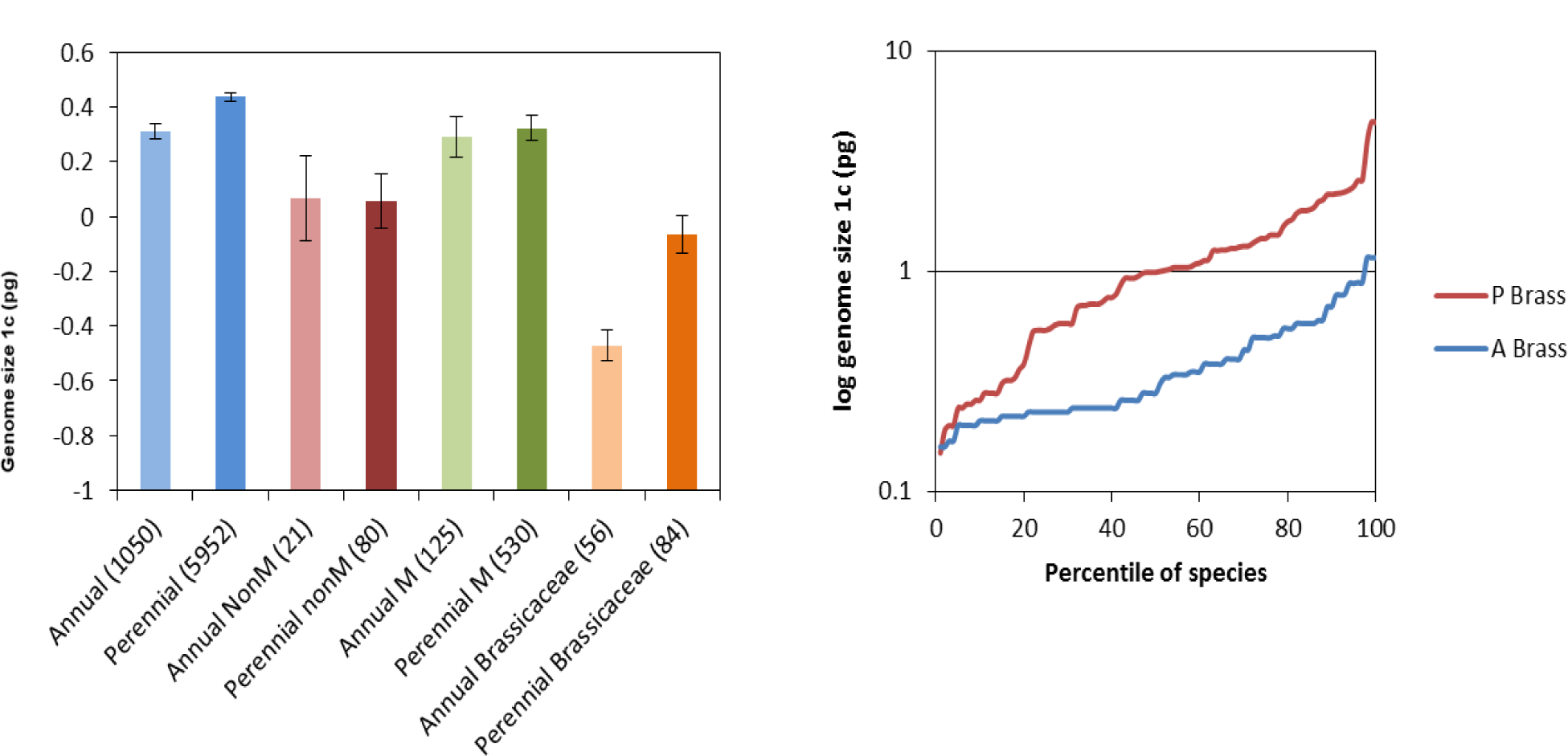
Log genome sizes comparisons between annuals and perennials. A. Comparisons between annuals and perennials in general (Annual, Perennial), plants with different relations to mycorrhiza (Annual NonM, Perennial NonM) and within the non-mycorrhizal family Brassicaceae (Annual Brassicaceae or A Brass, Perennial Brassicaceae or P Brass). Numbers in brackets represents number of plant species co-existing on lists for respective lifestyles and in the genome size database. Error bar represents 95% confidence intervals. Mean values with non-overlapping confidence intervals are significantly different (p=≪0.05). B. Sorted percentile plots from small genome size to high genome sizes for annual and perennial Brassicaceae species.

A multiplication of the whole genome as an increased ploidy from 2-4 should, although it could induce higher vigor, suddenly induce higher phosphorous drain from the nucleus on vital cell functions and thus a higher phosphorous demand for growth. Therefore, one can expect that a ploidy shift mainly happens for small nuclei since the stress on the phosphorous supply would be enormous and decrease plant fitness and lead to extinction even if it could be followed by a relatively fast reduction of DNA through evolution in the direction of removal of non-coding repeats (Šmarda et al., 2013). This should be reflected in an inverse relationship between n-size (genome/ploidy) and ploidy level as has been described earlier (Leitch, I. J. & Bennett, M. D., 2004). This I also found in my datasets. Even ploidies changes followed by changes in gene functions of multiplied genes followed by sexual recombination could result in better fitness and ultimately the formation of a new species (Leitch, I. J. & Bennett, M. D., 2004; Šmarda et al., 2013; Han et al., 2018). Thus, the punishment for a phosphorous drain by an increase in genome size should be smaller and the slope of the decrease in n-size less steep for even number ploidies than for odd number ploidies. This is also the case (**Fig. 3**), and suggests that gene diversification and speciation as well as P-stress induced relative fast genome shrinkages by a removal of repeats both relieves P-stress, weakening any potential correlation between speciation and genome size. Such a mechanism of fast removal of repeats connected with speciation due to gene duplications could explain why Kang et al. (Kang, Wang & Huang, 2015) found little phylogenetic signal correlated with genome size and P content in Karst plants while they found nitrogen content correlated with genome size and phylogeny. A better nutritional status for all nutrients except for P can be expected with evolution if P is always kept being limiting by the two mechanisms gene-duplication and accumulation of non-coding repeats.

**Figure 3.**
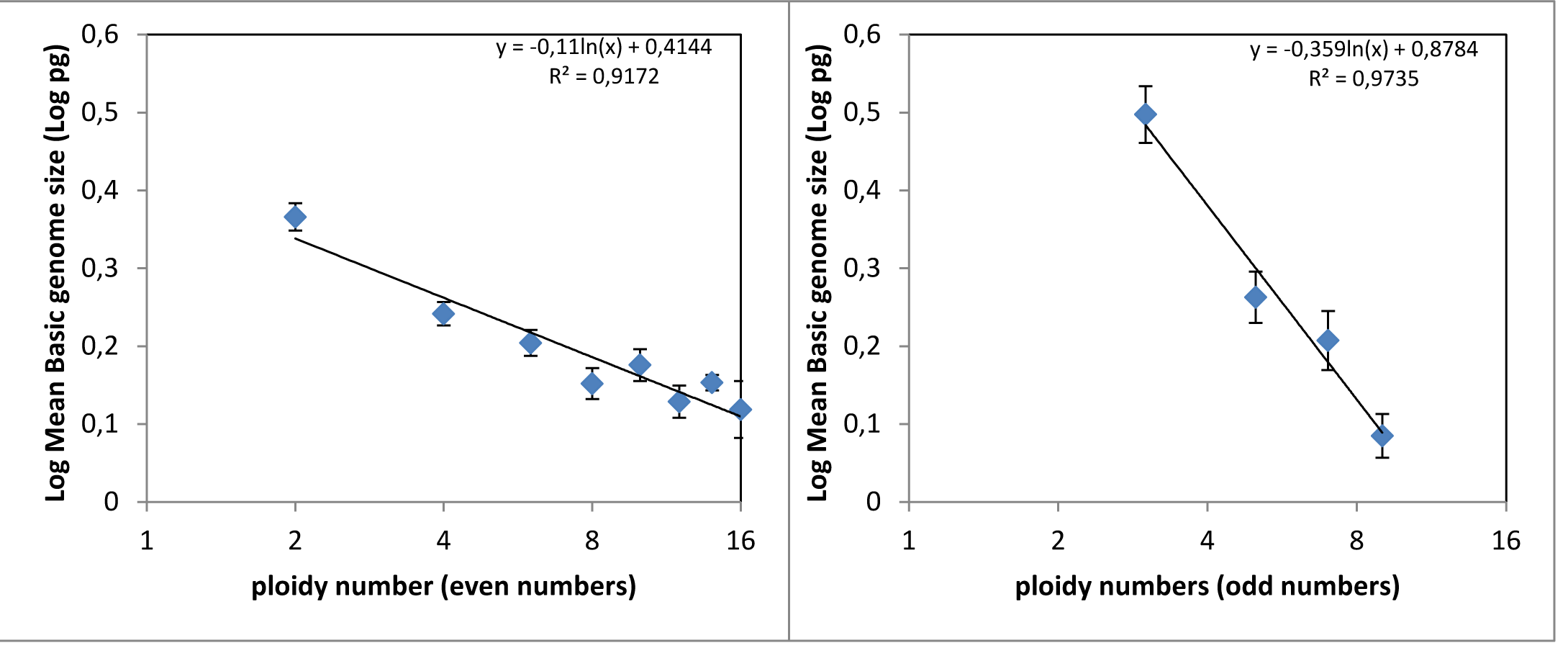
Log genome size versus log ploidy number. The basic haploid genome size decrease with higher ploidy number probably due to removal of non-coding repeats. As can be seen this removal is much weaker for plants with even ploidy numbers (left) than for odd ploidy numbers (right).

Because of the resulting P-stress of a whole genome duplication one can thus expect that a relaxed phosphorous limitation is needed to allow for ploidy shifts to higher ploidies. Since a perennial lifestyle seems to allow for larger genome sizes it should also allow for higher ploidy levels and there should thus be an overrepresentation of perennials among the plants with higher ploidies. If on the other hand higher ploidies precede development of a perennial lifestyle there should be an overrepresentation of higher ploidies among the annual plants. Similar reasoning can be made for ploidy and becoming mycorrhizal. I find a significant higher proportion (P_same_=0.04) (Fisher Exact test) of perennials to annuals (1493/213=6.46) among the higher ploidy plants (>2) than among all plants (4485/816=5.50) and especially compared to what is found among diploid (2n) plants (2992/585=5.11) (P_same_ =0.003). This is also obvious in the Brassicaceae family where the proportion of perennials to annuals in higher ploidy plants (>2) (38/13=2.92) was found to be significantly higher (P_same_ =0.044) than what was found in the whole family (69/47=1.47) and also significantly much higher (P_same_ =0.003) than found in the diploid brassicas (31/34=0.91). Thus developing a perennial lifestyle most likely precedes and allows for switching to higher ploidies. On the other hand I find no detectable significant relationships between mycorrhizal associations and ploidy level or mycorrhizal associations and perennial lifestyle in the data I have access to even if I find that the strictly non-mycorrhizal *Brassicaceae* contains significantly lower numbers (P_same_ =1.14E-06) of perennials to annuals (69/47=1.47) than mycorrhizal plants does (530/125=4.24). That could indicate that the mycorrhizal lifestyle favors perennial plants since there will then be no need to establish the interaction each year. It could also reflect that being mycorrhizal was the original lifestyle of land plants (Smith, Anderson & Smith, 2015). Thus, taken together with what is presented in Figure 2A, the extra benefit in phosphorous status by being mycorrhizal seems to offset any extra benefit in phosphorous status of becoming perennial but favors becoming perennial for other reasons.

### Conclusion

The available data suggests that P-availability is a driving factor in plant evolution due to the fact that the genomic DNA is a major P-pool that cannot be recycled for other uses (Figure 4). External environmental changes (like for example domestication), changes in lifestyle, mutations or mycorrhizal associations that improve P-nutrition, opens a temporary window for fast evolution of new traits through gene-duplication followed by evolution of new roles for the gene products. This window of opportunity is however soon closed by accumulation of repeats so that the system again becomes P-limited. The existence of this window of opportunity is also most likely why plants have not evolved more efficient mechanisms for the immediate removal of repeats since that would also remove genes that potentially can evolve into new uses. With severe P-limitation it seems to be possible to avoid accumulation of repeats or even to remove repeats which could explain why the carnivorous plants have very small genomes although they contain roughly the same number of functional genes as other plants (Leushkin et al., 2013; Fleischmann et al., 2014; Carretero-Paulet et al., 2015). All this above could also explain why there is a faster shrinking 1c genome size with higher ploidies for the odd ploidies compared to the even ploidies.

**Figure 4.**
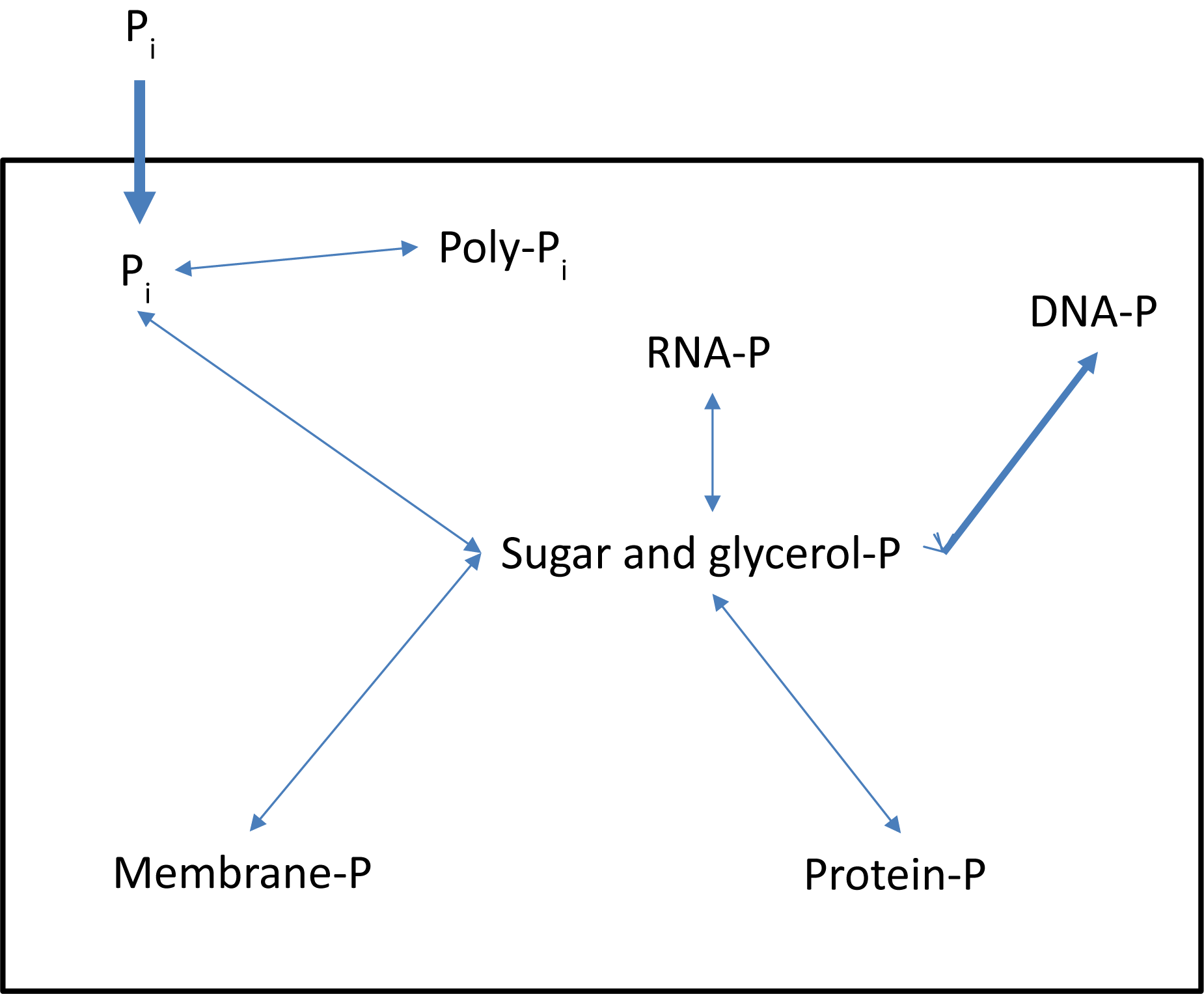
Schematic representation of the recycling and the main pools of phosphorous in a plant cell. Phosphorous taken up from the environment are distributed to different pools. DNA-P is the only large phosphorous pool in the cell that is a true sink and not possible to recycle P from for other uses without killing the cell.

### Dead end evolution

High P availability thus gives short term gains in competitive advantage (Pandit, White & Pocock, 2014) but drives plant evolution towards large nuclei with evolutionary difficulties to switch back to growth in low P niches. Large nuclei plants are short term winners but losers of tomorrow since they (the species) and their descendants are on a one way road towards extinction. The main reservoir for the evolution of the bulk of future plant species millions of years from now are most probably found in the plants that now have small nuclei.

## Methods

### Sources of data (See Supplementary file S1 *Web resources*)

Data on 1C genome angiosperm C-values was downloaded from the Kew database. Information on mycorrhizal, non-mycorrhizal and myco-heterotrophic lifestyle was mainly extracted from (Wang & Qiu, 2006). Information on parasitic and carnivorous lifestyles were downloaded and extracted.

### Data handling, data files and statistical programs used. (See Supplementary file S1 *Web resources*)

MS excel was used to handle, sort and plot the data as well as for the Fisher Exact test add-in. The most important handling was to determine which plant species where on the lists for different lifestyles and on 1C database list. All species occurring on both lists were included on the resulting lists so as not to introduce bias. The final Excel files used for the analysis are available in as supplementary data (**Supplementary Data S1-4**). The easy to use statistical freeware Past 3.18 was used for calculating upper and lower 95% confidence intervals using the BCa (adjusted percentile method) without assuming the distribution of the data that was not normally distributed and also for generating data for the percentile plots.

## Supporting information

List of Web resources

## Acknowledgements

I thank Professor Bjoern Hamberger, previous colleague at University of Copenhagen, now Michigan State University, U.S.A for triggering my curiosity in finding explanations for plant genome sizes by showing me the scientific paper about the very small carnivorous plants genomes (Leushkin et al., 2013) when we were teaching a course together and for further pointing out that plants species have a very large range of genome sizes without having many more genes, something I then was not aware of. Also, a special thanks to Professor Bjoern Hamberger for good and critical discussions in the beginning of this work and for encouraging me to gather data that could be used to test my hypotheses and publish. The basic support for this work comes from my employer Fujian Agriculture and University that gives me opportunity to freely explore different research topics. Finally, I want to thank the people and funders that keep the web-resources I have used available to anyone (See **Supplementary file S1** *Web resources*). Without that this work would not have been possible.

## Supplementary material

**File S1.** List of Web-resources

## Supplementary data files available at Figshare

**Data S1.** 1C genome size values, lifestyles and ploidy of Brassicaceae plants.

Private link: https://figshare.com/s/e247ace54392871e8ad5

DOI (reserved to be activated after manuscript acceptance): 10.6084/m9.figshare.8378414

**Data S2.** 1C genome size values, lifestyles and ploidy of carnivorous plants

Private link: https://figshare.com/s/dcdd06f6641c0b643bb5

DOI (reserved to be activated after manuscript acceptance): 10.6084/m9.figshare.8378474

**Data S3.** 1C genome size values, lifestyles and ploidy of mycorrhizal plants.

Private link: https://figshare.com/s/1f7dc82ddcc32489d9d4

DOI (reserved to be activated after manuscript acceptance): 10.6084/m9.figshare.8378477

**Data S4.** 1C genome size values, lifestyles and ploidy of parasitic plants.

Private link: https://figshare.com/s/0b8ed26c18b65d786fca

DOI (reserved to be activated after manuscript acceptance): 10.6084/m9.figshare.8378480

