## Supplementary material for "Reckless driving: Improved phosphorous availability is a most likely important driver and species killer in angiosperm evolution": List of Web resources

PLANT DATA

Angiosperm C-values and data on Non-perennial/Perennial lifestyle

<http://data.kew.org/cvalues/CvalServlet?querytype=2>

Parasitic plant genera

<https://www.aphis.usda.gov/plant_health/permits/organism/downloads/parasitic_plant_genera.pdf>

Carnivourus plants

<http://www.pinguicula.org/pages/pages_principales/SEZNAM%202004.pdf>

STATISTICS

Fisher Exact add-in for Excel

<http://www.obertfamily.com/software/fisherexact.html>

PAST statistical freeware

<https://folk.uio.no/ohammer/past/>
